# ORF3a mediated-incomplete autophagy facilitates SARS-CoV-2 replication

**DOI:** 10.1101/2020.11.12.380709

**Authors:** Yafei Qu, Xin Wang, Yunkai Zhu, Yuyan Wang, Xing Yang, Gaowei Hu, Chengrong Liu, Jingjiao Li, Shanhui Ren, Zixuan Xiao, Zhenshan Liu, Weili Wang, Ping Li, Rong Zhang, Qiming Liang

**Author notes:** These authors contributed equally (Yafei Qu, Xin Wang, Yunkai Zhu). Correspondence: Ping Li, Rong Zhang, Qiming Liang.

## Abstract

SARS-CoV-2 is the causative agent for the COVID-19 pandemic and there is an urgent need to understand the cellular response to SARS-CoV-2 infection. Beclin-1 is an essential scaffold autophagy protein that forms two distinct subcomplexes with modulators Atg14 and UVRAG, responsible for autophagosome formation and maturation, respectively. In the present study, we found that SARS-CoV-2 infection triggers an incomplete autophagy response, elevated autophagosome formation but impaired autophagosome maturation, and declined autophagy by genetic knockout of essential autophagic genes reduces SARS-CoV-2 replication efficiency. By screening 28 viral proteins of SARS-CoV-2, we demonstrated that expression of ORF3a alone is sufficient to induce incomplete autophagy. Mechanistically, SARS-CoV-2 ORF3a interacts with autophagy regulator UVRAG to facilitate Beclin-1-Vps34-Atg14 complex but selectively inhibit Beclin-1-Vps34-UVRAG complex. Interestingly, although SARS-CoV ORF3a shares 72.7% amino acid identity with the SARS-CoV-2 ORF3a, the former had no effect on cellular autophagy response. Thus, our findings provide the mechanistic evidence of possible takeover of host autophagy machinery by ORF3a to facilitate SARS-CoV-2 replication and raises the possibility of targeting the autophagic pathway for the treatment of COVID-19.

## MAIN TEXT

Severe Acute Respiratory Syndrome Coronavirus-2 (SARS-CoV-2) is a novel coronavirus confirmed as the causative agent of coronavirus disease 2019 (COVID-19) that began in Wuhan, China in late 2019 and spread worldwide^1,2^. As of 25 October 2020, there have been more than 42 million confirmed cases of SARS-CoV-2 infection with more than 1.1 million deaths attributed to the virus in 235 countries and territories. The clinical presentation of symptoms of COVID-19 ranges from asymptomatic to acute respiratory distress syndrome. The majority of cases will be mild to moderate but individuals with underlying comorbidities are at higher risk for developing more serious complications including respiratory failure, shock, and multiorgan system dysfunction^3^. While there continues to be unprecedented collaboration to facilitate the development of therapeutic neutralizing antibodies, vaccines, and small molecule drugs against COVID-19^4^, only one FDA-approved drug (Veklury [Remdesivir]) is currently available and there is an urgent need to better understand how SARS-CoV-2 manipulates the host responses.

SARS-CoV-2 is an enveloped, positive-sense, single-stranded RNA virus that engages human angiotensin-covering enzyme 2 (hACE2) to mediate host cell entry in the nasal passage, respiratory tract, and intestine^5^. Infection with SARS-CoV-2 can lead to excessive production of pro-inflammatory cytokines and dysregulation of type I interferon response^6–8^. Quantitative proteomic analysis of lung epithelial (A549) cells infected with SARS-CoV-2 and peripheral blood mononuclear cell (PBMC) specimens of COVID-19 patients revealed perturbations of a broad range of cellular signaling pathways, including macroautophagy^9^. Macroautophagy, hereafter referred to as autophagy, is a critical house-keeping process involving the formation of double-membrane autophagosomes that later fuse with lysosomes to degrade and recycle damaged organelles, unused proteins and invading pathogens^10^. In mammalian cells, this process is orchestrated by Beclin-1, a scaffolding protein that regulates the lipid kinase Vps34 (PI3KC3) and interacts with several cofactors/adaptor proteins (Atg14, UVRAG) to promote the formation of mutually exclusive Beclin-1-Vsp34 subcomplexes with distinct functions^10^. The Beclin-1-Vps34-Atg14 complex mainly positively regulates autophagy by promoting autophagosome formation while the Beclin-1-Vps34-UVRAG protein complex accelerates autophagosome maturation by promoting the fusion of autophagosomes with lysosomes to allow degradation and recycling of cellular components^11^.

Autophagy is an important homeostatic mechanism for cell protection and is evolutionarily conserved from yeast to higher eukaryotes; however a subset of viruses has subverted the autophagic pathway to benefit their replication^12^. For example, Zika virus utilizes NS4A and NS4B to inhibit the Akt-mTOR signaling pathway, leading to aberrant activation of autophagy and increased viral replication^13^; Kaposi’s sarcoma-associated herpesvirus vBcl2, vFLIP, and K7 proteins serve as anti-autophagy molecules to inhibit autophagosome formation or maturation via targeting Beclin-1, Atg3, and Rubicon, respectively^14–17^; and herpes simplex virus 1 encodes protein ICP34.5 which binds to Beclin-1 and inhibits its autophagy function^18^. Studies on other betacoronaviruses show that these viruses modulate autophagy but it is unknown whether viral evasion of autophagy is important in SARS-CoV-2 infection.

SARS-CoV-2 can infect multiple hACE2-expressing cell lines and in refractory cell lines after exogenous hACE2 expression; however, it is unclear if SARS-CoV-2 infection modulate cellular autophagy response. To evaluate autophagic activity upon SARS-CoV-2 infection, we measured microtubule-associated protein light chain 3 (LC3) conversion (soluble LC3-I to lipid bound LC3-II), which is correlated with the formation of autophagosomes and is widely used as a marker to monitor autophagy^19^. However, LC3-II is degraded together with the contents of the autophagosome and increased LC3-II may reflect either increased autophagosome formation or, alternatively, decreased autolysosome degradation (i.e. due to impairment of fusion between autophagosomes and lysosomes in autophagy). Thus, it is important to measure LC3-II turnover both in the presence and absence of autophagy inhibitors, commonly achieved using bafilomycin A1, a macrolide antibiotic that prevents maturation of autophagic vacuoles by inhibiting autophagosome-lysosome fusion^19^. In stable hACE2 expressing HeLa cells infected with SARS-CoV-2 strain SH01, we detected dramatic conversion of LC3-I to LC3-II in both the presence and absence of the lysosome inhibitor bafilomycin A1 (**Fig. 1a**), indicating that infection caused the upregulation of autophagosome formation. Next, we measured ubiquitin receptor SQSTM1/p62 turnover, the levels of which usually correlates with autophagic flux^19^. SQSTM1/p62 did not decline during the course of SARS-CoV-2 infection (**Fig. 1a**), which strengthens the interpretation that infection blocked autophagosome-lysosome fusion and infected cells cannot form autolysosomes that degrade or recycle their contents. Similar LC3 conversion and SQSTM1/p62 stability were also observed in infected Calu-3 and Vero-E6 cell lines (**Fig. S1a-b**), suggesting SARS-CoV-2 induces incomplete autophagy in infected cells, which is consistent with speculations that SARS-CoV-2 infection suppresses autophagic flux^20^.

**Figure 1:**
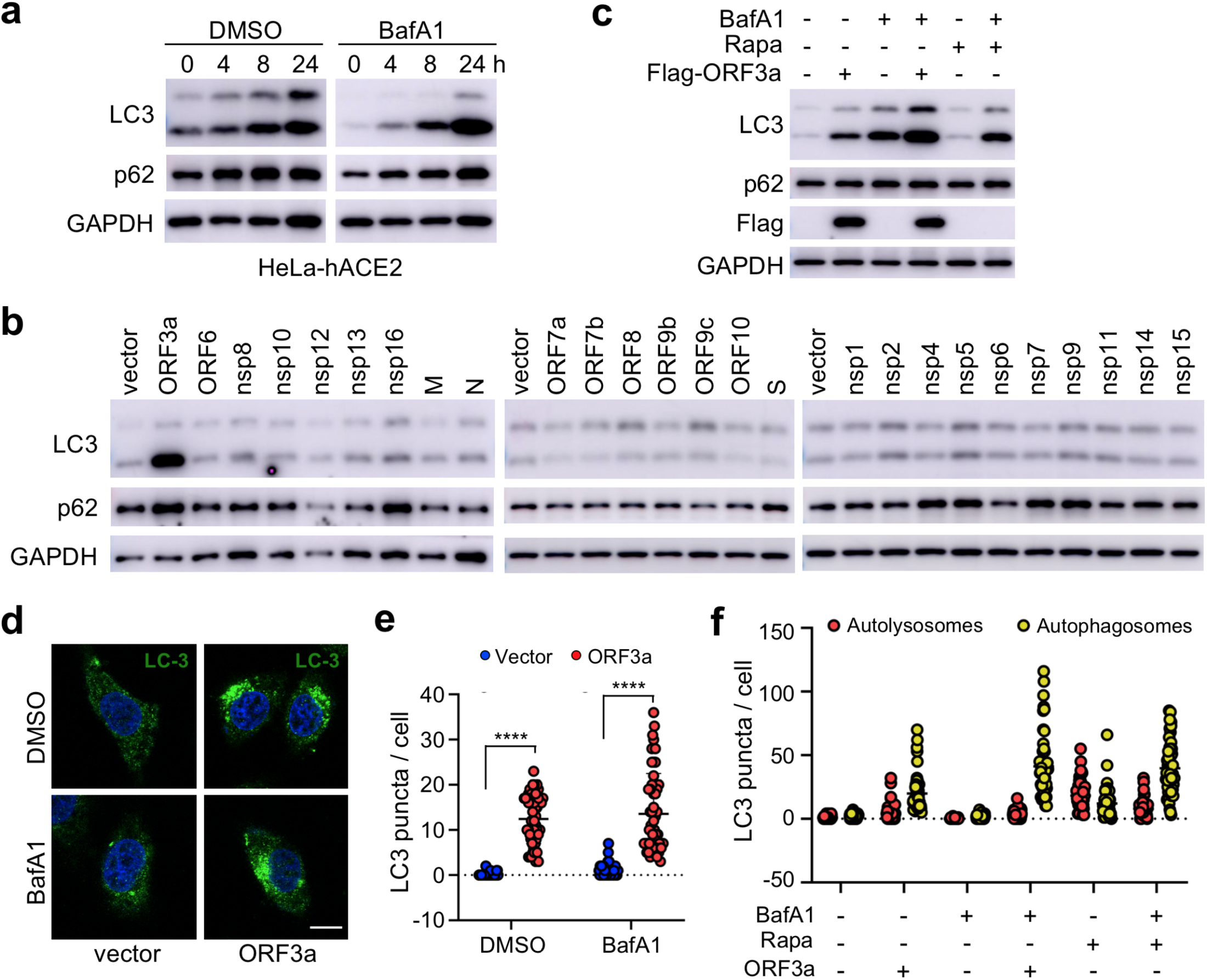
SARS-CoV-2 ORF3a triggers incomplete autophagy. (**a**) SARS-CoV-2 infection triggers incomplete autophagy. HeLa-hACE2 cells were infected with SARS-CoV-2 (MOI = 1) in the presence or absence of bafilomycin A1 (BafA1, 100nM) and cell lysates were collected at indicated time point for immunoblotting (IB) with indicated antibodies. (**b**) Screening of SARS-CoV-2-encoded proteins for LC3-I to LC3-II conversion in Hela cells. (**c**) SARS-CoV-2 ORF3a induces incomplete autophagy. Hela-vector or Hela-ORF3a cell lines were treated with indicated drugs (BafA1, 100nM; rapamycin, 1μM) for 4h and the cell lysates were collected for IB with indicated antibodies. (**d-e**) SARS-CoV-2 ORF3a induces LC3 puncta formation. Hela-vector or Hela-ORF3a cell line were treated with BafA1 (100nM) and endogenous LC3 puncta were immunostained (**d**) and quantified (**e**). Scale bar, 15μm. Mean ± SEM; n=50; \*\*\*\**p*<0.0001 by Student’s t test. (**f**) Hela-vector or Hela-ORF3a cell line were transfected with ptf-LC3 and transfected cells were treated with indicated drugs (BafA1, 100nM; rapamycin, 1μM) for 4h. Cells were fixed and the LC3 puncta were quantified as indicated. Scale bar, 15μm.

Since SARS-CoV-2 infection triggers an incomplete autophagy response in multiple cell lines, we attempted to determined which viral components can modulate the cellular pathways that control this process. The RNA genome of SARS-CoV-2 encodes 28 viral proteins, including 16 non-structural proteins (nsp1-16), 4 structural proteins [glycoprotein spike (S), membrane (M), envelope (E), and nucleocapsid (N)], and 8 accessory proteins (ORF3a, ORF6, ORF7a, ORF7b, ORF8, ORF9b, ORF9c, and ORF10). We screened each individual SARS-CoV-2 gene for LC3 conversion in Hela cells and found that ORF3a expression resulted in a significant increase in the conversion of LC3-I to LC3-II (**Fig. 1b**). However, other viral proteins had little or no effect (**Fig. 1b**). Similar findings were recapitulated when ORF3a was expressed in Hela, Vero-E6, and Calu-3 cells in the presence and absence of bafilomycin A1 (**Fig. 1c** and **S1c-d**), suggesting ORF3a itself can efficiently induce autophagosome accumulation. We also observed that SARS-CoV-2 ORF3a expression did not trigger the degradation of SQSTM/p62 in above cell lines (**Fig. 1c**), indicating the fusion step between autophagosome and lysosome (autophagosome maturation) is blocked. Consistently, expression of ORF3a led to dramatically elevation of LC3 puncta per cell in Hela cells with or without Bafilomycin A1 treatment (**Fig. 1d-e**), confirming the efficient induction of autophagosome formation. In addition, we utilized tandem fluorescent-tagged LC3 (GFP-mRFP-LC3) to differentiate between autophagosomes and autolysosomes upon SARS-CoV-2 ORF3a expression. Since GFP is unstable in the acidic environment of lysosome, unmatured autophagosomes and matured autolysosome are labelled by yellow LC3 puncta (GFP+mRFP) or red LC3 puncta (mRFP), respectively^21^. In Hela-vector control cells, rapamycin-triggered autophagy resulted in the increase of both yellow and red puncta, while Bafilomycin A1 treatment only generated yellow autophagosome LC3 puncta by blocking autophagosome-lysosome fusion (**Fig. 1f** and **S1e**). Similar to treatment with both rapamycin and Bafilomycin A1, SARS-CoV-2 ORF3a significantly increased unmatured autophagosome (yellow puncta) but had little effect on matured autolysosome (red puncta) (**Fig. 1f** and **S1e**), indicating that ORF3a triggers autophagy but inhibits the fusion between autophagosomes and lysosomes. Taken together, our findings suggest that ORF3a efficiently triggers incomplete autophagy in multiple cell lines, which phenocopies SARS-CoV-2 infection.

Next, we sought to determine the mechanism through which SARS-CoV-2 ORF3a induces incomplete autophagy. In recent months, four independent virus-host interactomes analyzed by network based-approach provided insights into viral pathogenesis^9,22–24^, but it remains unclear how SARS-CoV-2 interacts with the cellular autophagy machinery to trigger incomplete autophagy. Using yeast two-hybrid screening with SARS-CoV-2 ORF3a as a bait, we captured a novel protein-protein interaction with UVRAG (UV resistance-associated gene), a key autophagy regulator that activates Beclin-1-Vps34 complex to promote autophagosome maturation and suppresses the proliferation of human colon cancer cells^25,26^. SARS-CoV-2 ORF3a interacted with UVRAG in yeast (**Fig. 2a**). Among Beclin-1 complex components, UVRAG showed the strongest binding affinity to ORF3a in HEK293T cells (**Fig. 2b**). Detailed mapping suggested that ORF3a interacted with the N-terminal C2 and coiled coil domain (CCD) of UVRAG (**Fig. S2a-b**). The CCD provides a platform for many important protein interactions and is required for UVRAG binding to Beclin-1^25^. ORF3a also bond to and colocalized with endogenous UVRAG in Hela cells (**Fig. 2c** and **S2c**), further confirming UVRAG is the potential host target for SARS-CoV-2 ORF3a. Beclin-1 is an essential autophagy protein that orchestrates the assembly of functionally distinct Beclin-1-Vps34 multiprotein complexes that regulate autophagosome formation and maturation^11^. The core complex consists of Beclin-1, Vps34, and Vps15 which interacts with key modulators Atg14 and UVRAG to form mutually exclusive Atg14- or UVRAG-containing Beclin-1-Vps34 subcomplexes that positively regulates autophagy by promoting early-stage autophagosome formation and late-stage autophagosome maturation, respectively^11^. Since ORF3a binds to UVRAG and triggers incomplete autophagy, we speculated that ORF3a expression may lead to aberrant redistribution of Atg14- or UVRAG-containing Beclin-1-Vps34 complexes and disrupt the balance between autophagosome formation and maturation. To test our hypothesis, we purified the Beclin-1 complexes, UVRAG complexes, and Atg14 complexes, respectively, in the presence and absence of SARS-CoV-2 ORF3a. In line with previous reports, Atg14 and UVRAG were not present in the same complex and Rubicon is only associated with Beclin-1-Vps34-UVRAG complexes in the absence of ORF3a^27,28^ (**Fig. 2d-e** and **S2d**). In contrast, the presence of ORF3a significantly reduced the interaction between UVRAG and Beclin-1 or other Beclin-1 complex components, such as Vps34 and Rubicon, suggesting that Beclin-1-Vps34-UVRAG complex formation is disrupted by SARS-CoV-2 ORF3a (**Fig. 2d-e**). Furthermore, SARS-CoV-2 ORF3a expression enhanced the interaction between Atg14 and Beclin-1, suggesting dissociation between Beclin-1 and UVRAG facilitated Beclin-1-Vps34-Atg14 complex assembly (**Fig. 2d** and **S2d**). Taken together, these findings demonstrate that the ORF3a-UVRAG interaction renders UVRAG less competitive than Atg14 for Beclin-1 binding and shifts the balance between Beclin-1-Vps34-Atg14 and Beclin-1-Vps34-UVRAG complex assembly, resulting in more efficient autophagosome formation but inhibition of autophagosome maturation.

**Figure 2.**
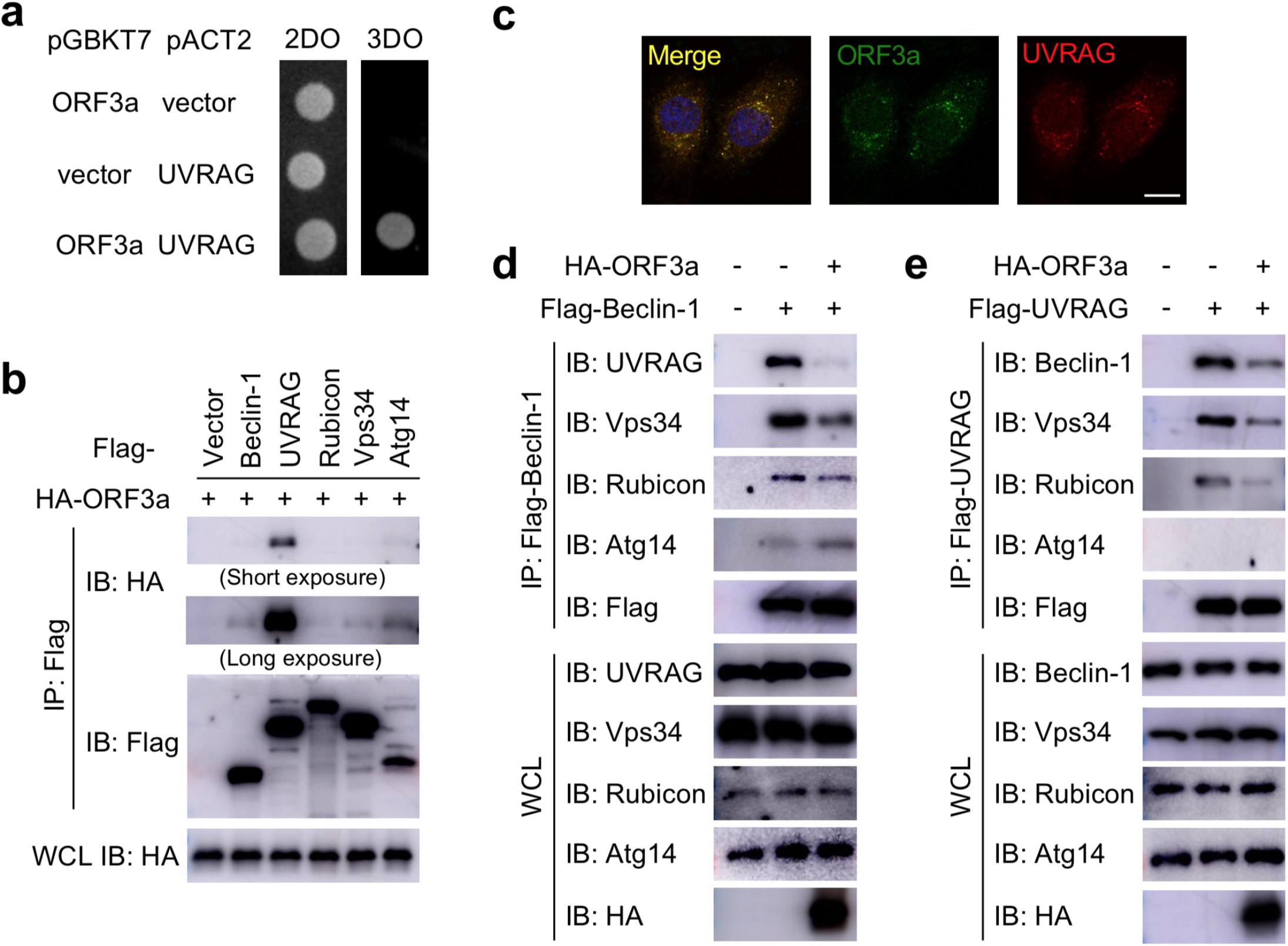
SARS-CoV-2 ORF3a targets UVRAG to modulate Beclin-1 complex formation. (**a**) SARS-CoV-2 ORF3a interacts with UVRAG in yeast two-hybrid system. (**b**) SARS-CoV-2 ORF3a binds to UVRAG in HEK293T cells. HEK293T cells were co-transfected with indicated plasmids and cell lysates were collected and subjected to immunoprecipitation (IP) and IB with indicated antibodies at 48h post-transfection. (**c**) SARS-CoV-2 ORF3a co-localizes with endogenous UVRAG. Hela-ORF3a stable cells were subjected to immunostaining with antibodies against Flag or UVRAG. Scale bar, 15μm. (**d-e**) SARS-CoV-2 ORF3a blocks Beclin-1-Vps34-UVRAG complex formation. HEK293T cells were co-transfected with indicated plasmids and cell lysates were collected and subjected to IP and IB with indicated antibodies at 48h post-transfection.

The genome of SARS-CoV-2 is closely related to SARS-CoV, the first deadly coronavirus that caused the SARS epidemic in 2003^29^. Since ORF3a^SARS^ shares 72.7% amino acid sequence identity to ORF3a^SARS-2^ (**Fig. 3a**), we sought to determine whether the ORF3a homologue from SARS-CoV could trigger similar incomplete autophagy. We generated a stable Hela cell line that expresses ORF3a^SARS^ and examined its effects on autophagy markers, including LC3-I to LC3-II conversion and LC3 puncta formation. Unlike ORF3a^SARS-2^, ORF3a^SARS^ expression led to neither dramatic increase in the amount of LC3-II nor accumulation of LC3 puncta in the presence or absence of Bafilomycin A1 (**Fig. 3b-c** and **S3a**), indicating ORF3a^SARS^ cannot efficiently induce the formation of autophagosomes. For tandem fluorescent-tagged LC3 (GFP-mRFP-LC3) assay, ORF3a^SARS^ expression had little effects on autophagosome (yellow puncta) and autolysosome (red puncta), while ORF3a^SARS-2^ significantly enhanced autophagosome formation but did not affect autolysosomes, when compared to vector control, further confirming that ORF3a^SARS-2^ but not ORF3a^SARS^ modulates cellular autophagy response (**Fig. S3b-c**). Furthermore, ORF3a^SARS^ did not interact with endogenous UVRAG (**Fig. 3d**) and could not modulate the formation of Beclin-1-Vps34-UVRAG or Beclin-1-Vps34-Atg14 complexes (**Fig. S3d-f**). These results show that ORF3a-mediated modulation of autophagy response is a unique signature of SARS-CoV-2 infection and may play an important role in controlling viral replication and pathogenesis.

**Figure 3:**
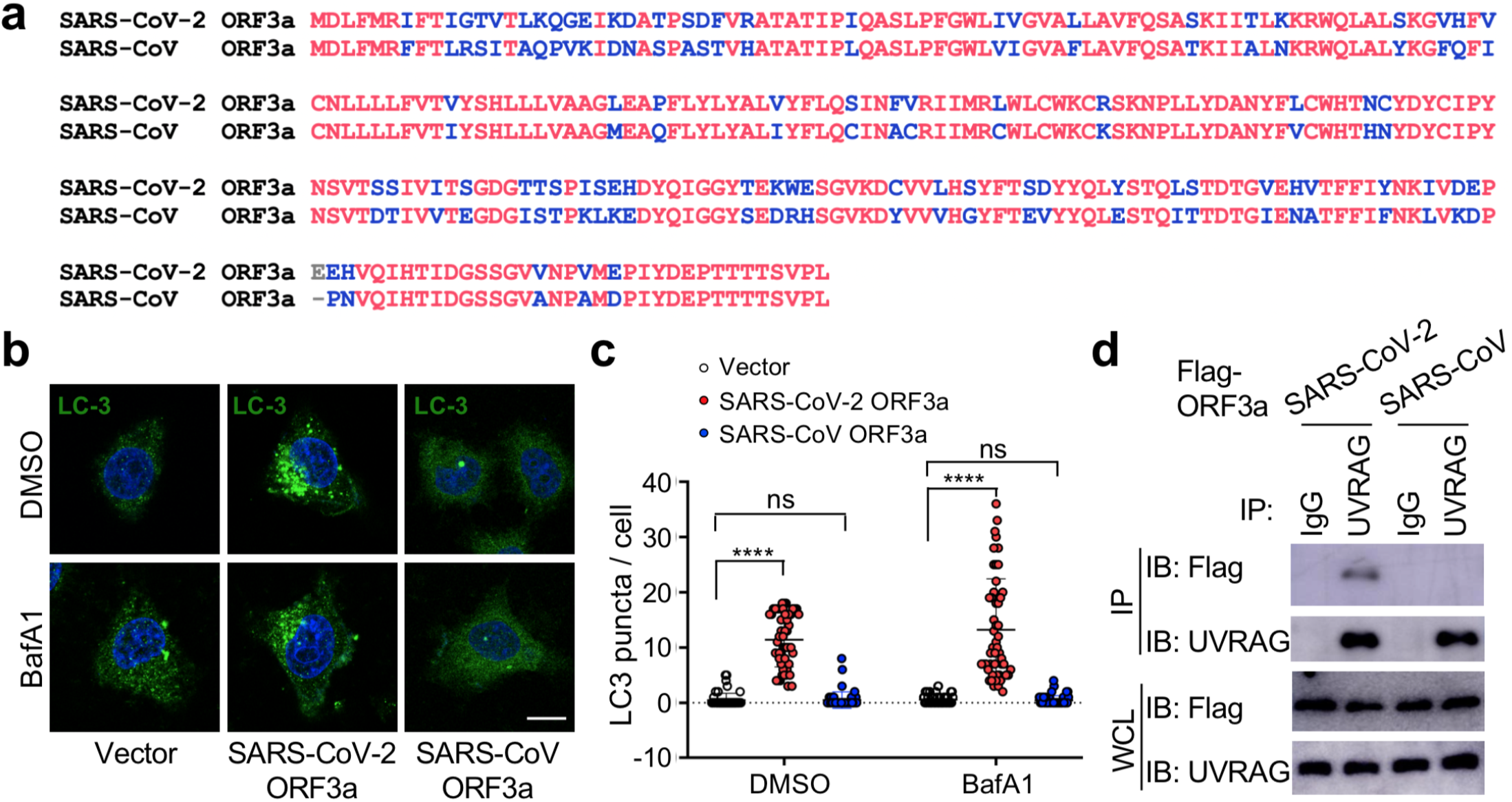
SARS-CoV ORF3a cannot induce autophagy. (**a**) Sequence alignment between SARS-CoV-2 ORF3a and SARS-CoV ORF3a. (**b-c**) SARS-CoV ORF3a does not affect autophagy. Hela-vector, Hela-ORF3a^SARS2^, or Hela-ORF3a^SARS^ cell line were treated with BafA1 (100nM) and endogenous LC3 puncta were immunostained (**b**) and quantified (**c**). Scale bar, 15μm. Mean ± SEM; n=50; *ns* and \*\*\*\**p*<0.0001 by Student’s t test. (**d**) ORF3a^SARS-2^ but not ORF3a^SARS^ interacts with endogenous UVRAG. HEK293T cells were transfected with Flag-ORF3a^SARS^ or Flag-ORF3a^SARS-2^ and cell lysates were collected and subjected to IP and IB with indicated antibodies at 48h post-transfection.

Finally, we analyzed the effect of autophagy on SARS-CoV-2 replication using cell lines with genetic abrogation of autophagic essential genes. Atg3 and Atg5 are two indispensable proteins that mediate vesicle elongation during the autophagosome formation process and genetic knockout of Atg3 or Atg5 blocks LC3 conversion and SQSTM1/p62 degradation in MEFs^10^ (**Fig. 4a**). Using MEF stable expressing hACE2 receptor, we investigated how abrogation of cellular autophagy response by Atg3 or Atg5 knockout affect SARS-CoV-2 replication. As shown in Figure 4b-e, the autophagy machinery is required for efficient SARS-CoV-2 replication, and infection of autophagy-deficient cells resulted a significant reduction in viral yield compared to yield in control cells, as shown by immunoblot analyses of viral N protein expression (**Fig. 4b-c**) and quantitative RT-PCR of viral transcripts (**Fig. 4d-e**). Collectively, these results demonstrated that cellular autophagy response is required for efficient SARS-CoV-2 replication.

**Figure 4.**
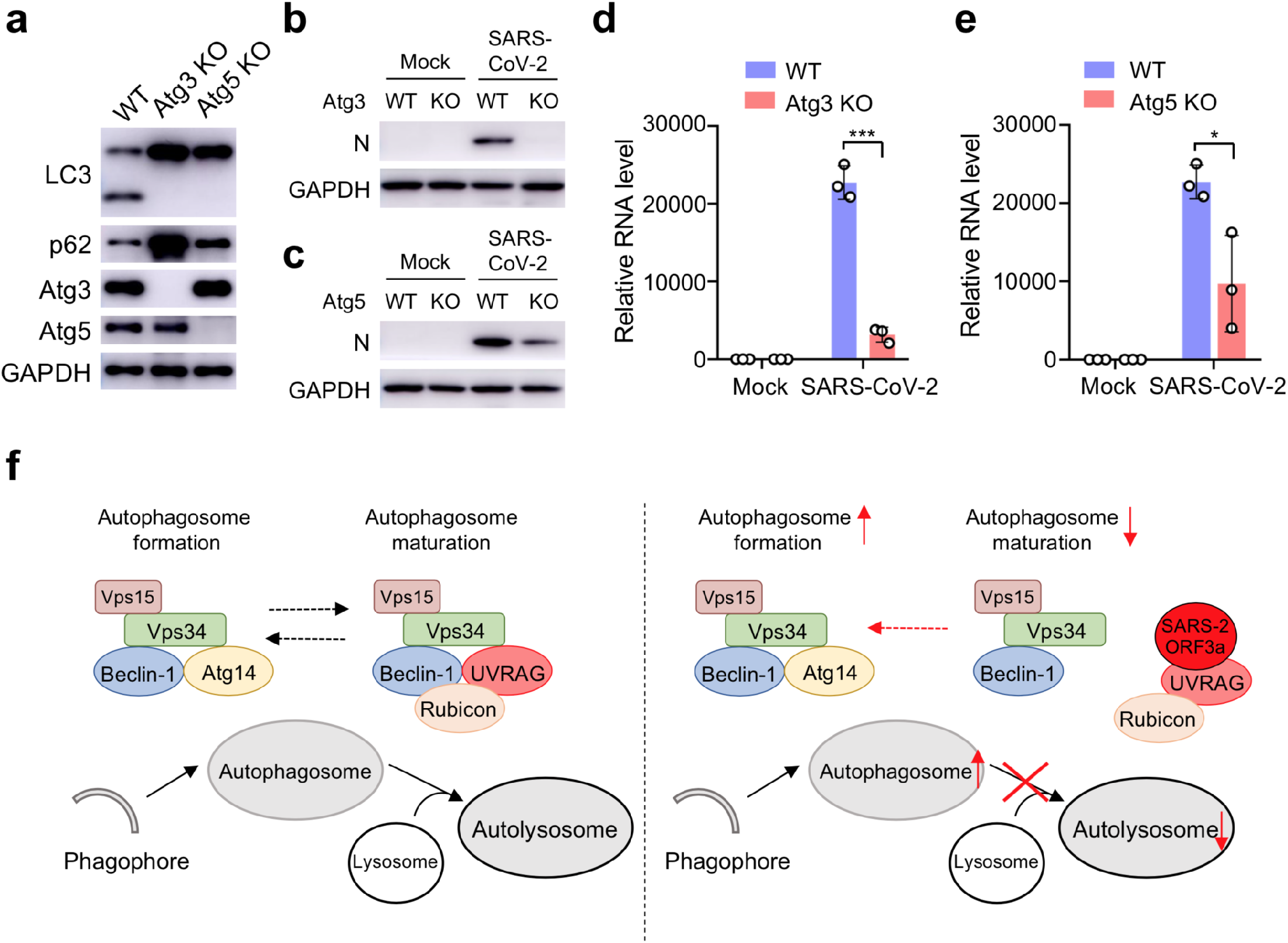
Cellular autophagy is required for efficient SARS-CoV-2 replication. (**a**) Genetic abrogation of Atg3 or Atg5 blocks autophagy response in MEF-hACE2 cell lines. (**b-e**) Atg3 or Atg5 KO reduces SARS-CoV-2 replication efficiency in MEF-hACE2 stable cells. MEF-hACE2^WT^, MEF-hACE2^Atg3 KO^, or MEF-hACE2^Atg5 KO^ were infected with SARS-CoV-2 (MOI = 1) and virus replication was examined by IB or viral N protein (**b-c**) or quantitative RT-PCR of viral transcripts (**d-e**). Mean ± SEM; n=3; \**p*<0.05 and \*\*\**p*<0.001 by Student’s t test. (**f**) Schematic diagram of SARS-CoV-2 ORF3a-mediated incomplete autophagy.

Many coronaviruses trigger autophagy in infected cells^30,31^, however, how SARS-CoV-2 modulates cellular autophagy and whether autophagy affects SARS-CoV-2 replication remain elusive. In this study, we found that ORF3a triggers incomplete cellular autophagy to generate double-membrane autophagosome vesicles, which are required for efficient SARS-CoV-2 replication. Mechanistically, SARS-CoV-2 ORF3a interacts with UVRAG to positively regulate Beclin-1-Vps34-Atg14 complex formation but suppress the Beclin-1-Vps34-UVRAG complexes, thus elevating autophagosome formation and blocking autophagosome maturation (**Fig. 4f**). Given autophagy’s role in the innate antiviral immune response to restrict infecting pathogens, it is not surprising that viruses are in a constant arms race to remodel the autophagic membranes for their own benefit during replication^12^. SARS-CoV-2 ORF3a hijacks the autophagy machinery to generate double-membrane autophagosome vesicles to facilitate viral replication but arrests the autophagosomes prior to lysosome fusion to avoid succumbing to lysosomal degradation. Interestingly, although ORF3a^SARS^ shares 72.7% amino acid identity with the ORF3a^SARS-2^, the former had no effect on the formation of double-membrane autophagosome vesicles. Coronaviruses are known to rely on the formation of convoluted autophagosome-like double membrane vesicles for optimal replication^30,32^, and ORF3a-mediated autophagosome formation may be one source of these double-membrane replication organelles. Veklury (Remdesivir) has just been approved for the treatment of SARS-CoV-2 infection and several preclinical investigations repurposing several FDA-approved drugs are underway. Since elevated autophagy facilitates coronaviruses replication efficiency^30^, the drugs that enhance autophagy should be avoided throughout the course of treatment. Antimalarial drugs chloroquine (CQ) and hydroxychloroquine are lysosomotropic agents that inhibit the pH-dependent replication steps in several viruses and many publications have urged the use of CQ as a potential treatment for COVID-19 based on *in vitro* considerations^33–37^. However, there are still discrepancies and concerns on the sensitivity and therapeutic range of CQ as its effectiveness on limiting SARS-CoV-2 replication did not extend to TMPRSS2-expressing human lung cells^38^, suggesting the cell specific responses may exist. In addition, CQ-mediated inhibition of cellular autophagy response may also contribute to its effects on SARS-CoV-2^19^. In summary, our work highlights the mechanism of how SARS-CoV-2 co-opts the autophagy pathway to enhance its own replication and spread, and raises the possibility to targeting the autophagic pathway for the treatment of COVID-19.

## AUTHOR CONTRIBUTION

Q.L. conceived of the research, designed the study, and wrote the manuscript. Y.Q., X.W., X.Y., C.L., J.L., Z.L., W.W., and P.L. performed the experiments and analyzed data. R.Z., Y.Z., Y.W., G.H., and S.R. performed the SARS-CoV-2 infection experiments in BSL-3 Laboratory. Z.X. helped with visual representation of data and edited the manuscript. All authors commented on the manuscript.

## ACKNOWLEDGMENTS

We thank Drs. Peihui Wang (Shandong University, China) and Qing Zhong (Shanghai Jiao Tong University, China) for providing SARS-CoV ORF3a plasmid and MEFs (Atg3 KO and Atg5 KO), respectively. This work was supported by grants from National Key Research and Development Project of China (2018YFA0900802 and 2020YFA0707701), National Natural Science Foundation of China (31770176 and 32041005), the Program for Professor of Special Appointment (Eastern Scholar) at Shanghai Institutions of Higher Learning, Shanghai Science and Technology Commission (20YF1442500, 20431900401), Shanghai Municipal Health Commission (2018YQ40, 201940179, and 20204Y0347), Innovative Research Team of High-level Local Universities in Shanghai, the Interdisciplinary Program of Shanghai Jiao Tong University (YG2020YQ14), and the SII Challenge Fund for COVID-19 Research. The Core Facility of Basic Medical Sciences, Shanghai Jiao Tong University School of Medicine provided the cDNA plasmids for cloning. We wish to acknowledge Prof. Di Qu, Dr. Xia Cai, and other colleagues at the Biosafety Level 3 Laboratory of Fudan University for help with experiment design and technical assistance.

## DECLARATION OF INTERESTS

The authors declare no competing interests.

## Methods

### Viruses, Plasmid construction, and Cell culture

SARS-CoV-2 strain SH01 (GenBank: MT121215.1) was provided by Dr. Rong Zhang (Fudan University, China). SARS-CoV-2 stocks were propagated in Vero-E6 cells and the titer of SARS-CoV-2 stocks were determined by standard plaque assay on Vero-E6 cells as described previously^39^. Experiments related to SARS-CoV-2 infection are performed in BSL-3 laboratory in Fudan University.

HEK293T (ATCC, #CRL-11268), Hela (ATCC, #CCL-2), A549 (ATCC, #CCL-185), and MEF (wild-type, Atg3 KO, or Atg5 KO, gift from Dr. Qing Zhong at Shanghai Jiao Tong University School of Medicine, China) cells were maintained in Dulbecco’s modified Eagle’s medium (DMEM; Gibco-BRL) containing 4 mM glutamine and 10% FBS. Vero-E6 cells (gift from Dr. Gang Long at Institut Pasteur of Shanghai, Chinese Academy of Sciences) were cultured in DMEM with 5% FBS and 4 mM glutamine. Calu-3 cells (gift from Dr. Rong Zhang at Fudan University, China) were cultured in DMEM with 20% FBS and 4 mM glutamine. Transient transfections were performed with Lipofectamine 3000 (Thermo Fisher Scientific, #3000015).

SARS-CoV-2 genes in pLVX-3xFlag-MCS-P2A-tagRFP (puro) have been described previously^22^. ORF3a^SARS-2^ were subcloned into pEF-MCS-3xHA, pCDH (Hygro), or pGBTK7 vectors. The ORF3a^SARS^ template was kindly provided by Dr. Peihui Wang (Shandong University, China) and cloned into pLVX vector. The constructs encoding Beclin-1, Atg14, Vsp34, UVRAG, and Rubicon have been previously described^17^. All constructs were sequenced using an ABI PRISM 377 automatic DNA sequencer to verify 100% correspondence with the original sequence.

### Detection of Autophagy

For the LC3 mobility shift assay, SARS-CoV-2-infected or ORF3a expressing cells were washed with cold PBS, lysed with 1% Triton X-100, and then subjected to immunoblot analysis (15% SDS-PAGE) with antibodies against LC3 (Cell Signaling, #3868) or SQSTM1/p62 (Cell Signaling, #39749). LC3-I is about 14 kD, and lipidated LC3 (LC-II) is about 16 kD. The ratio of LC3-II to LC3-I represents autophagy level. Autophagy was also assessed by LC3 redistribution. SARS-CoV-2-infected or ORF3a expressing cells were fixed and endogenous LC3 was detected by confocal microscope. To quantitate LC3-positive autophagosomes per cell, cells from five random fields (>50 cells) were counted. The number of LC3 puncta per cells was determined by NIH ImageJ software in three independent experiments.

### Immunoblot analysis

Cell lysates were collected in 1% Triton X-100 buffer and quantified by Bradford protein assay (Thermo Scientific). Proteins were separated by SDS-PAGE and transferred to PVDF membrane (Bio-Rad) by semi-dry transfer at 25V for 30 minutes. All membranes were blocked in 5% milk in PBST and probed overnight with indicated antibodies in 5% BSA at 4°C. Primary antibodies included: mouse Flag (Sigma, #F1804), rat Flag-HRP (Biolegend, #637311), mouse HA (BioLegend, #901515), rabbit SQSTM1/p62 (Cell Signaling, #39749), rabbit LC3 (Cell Signaling, #3868), rabbit Beclin-1 (Cell Signaling, #3495), rabbit UVRAG (Cell Signaling, #13115), rabbit Atg14 (Cell Signaling, #96752), rabbit Rubicon (Cell Signaling, #8465), rabbit Vps34 (Cell Signaling, #4263), rabbit Atg3 (Cell Signaling, #3415), rabbit Atg5 (Cell Signaling, #12994), mouse SARS-CoV-2 N (GeneTex, #GTX635689), and mouse GAPDH (Santa Cruz, #365062). Appropriate HRP-conjugated secondary antibodies were incubated on membranes in 5% milk and bands were developed with ECL reagent (Thermo Scientific) and imaged on a Fuji LAS-4000 imager.

### RNA extraction and quantitative RT-PCR

Total RNA was isolated from cells with the RNeasy Mini Kit (Qiagen, #74106) and treated with RNase-free DNase according to the manufacturer’s protocol. All gene transcripts were quantified by quantitative PCR using qScript™ One-Step qRT-PCR Kit (Quanta Biosciences, #95057-050) on CFX96 real-time PCR system (Bio-Rad). Primer sequences for qPCR were as follow: SARS-CoV-2-N forward: GACCCCAAAATCAGCGAAAT, SARS-CoV-2-N reverse: TCTGGTTACTGCCAGTTGAATCTG; 18S forward: GTAACCCGTTGAACCCCATT, 18S reverse: CCATCCAATCGGTAGTAGCG.

### Quantification and Statistical Analysis

All data were expressed as Mean ± SEM as indicated. Statistical significance across two groups was tested by Student’s t-test. *P*-values of less than 0.05 were considered significant.

## Notes

### Competing Interest Statement

The authors have declared no competing interest.

## REFERENCE

1. Wu, F. et al. A new coronavirus associated with human respiratory disease in China. Nature 579, 265–269 (2020).

2. Zhou, P. et al. A pneumonia outbreak associated with a new coronavirus of probable bat origin. Nature 579, 270–273 (2020).

3. Huang, C. et al. Clinical features of patients infected with 2019 novel coronavirus in Wuhan, China. Lancet Lond. Engl. 395, 497–506 (2020).

4. Florindo, H. F. et al. Immune-mediated approaches against COVID-19. Nat. Nanotechnol. 1–16 (2020) doi:10.1038/s41565-020-0732-3.

5. Hoffmann, M. et al. SARS-CoV-2 Cell Entry Depends on ACE2 and TMPRSS2 and Is Blocked by a Clinically Proven Protease Inhibitor. Cell 181, 271–280.e8 (2020).

6. Blanco-Melo, D. et al. Imbalanced Host Response to SARS-CoV-2 Drives Development of COVID-19. Cell 181, 1036–1045.e9 (2020).

7. Hadjadj, J. et al. Impaired type I interferon activity and inflammatory responses in severe COVID-19 patients. Science (2020) doi:10.1126/science.abc6027.

8. Zhang, X. et al. Viral and host factors related to the clinical outcome of COVID-19. Nature 583, 437–440 (2020).

9. Stukalov, A. et al. Multi-level proteomics reveals host-perturbation strategies of SARS-CoV-2 and SARS-CoV. bioRxiv 2020.06.17.156455 (2020) doi:10.1101/2020.06.17.156455.

10. Dikic, I. & Elazar, Z. Mechanism and medical implications of mammalian autophagy. Nat. Rev. Mol. Cell Biol. 19, 349–364 (2018).

11. Kang, R., Zeh, H. J., Lotze, M. T. & Tang, D. The Beclin 1 network regulates autophagy and apoptosis. Cell Death Differ. 18, 571–580 (2011).

12. Choi, Y., Bowman, J. W. & Jung, J. U. Autophagy during viral infection - a double-edged sword. Nat. Rev. Microbiol. 16, 341–354 (2018).

13. Liang, Q. et al. Zika Virus NS4A and NS4B Proteins Deregulate Akt-mTOR Signaling in Human Fetal Neural Stem Cells to Inhibit Neurogenesis and Induce Autophagy. Cell Stem Cell 19, 663–671 (2016).

14. Gao, H., Song, Y., Liu, C. & Liang, Q. KSHV strategies for host dsDNA sensing machinery. Virol. Sin. 31, 466–471 (2016).

15. Lee, J.-S. et al. FLIP-mediated autophagy regulation in cell death control. Nat. Cell Biol. 11, 1355–1362 (2009).

16. Liang, C., E, X. & Jung, J. U. Downregulation of autophagy by herpesvirus Bcl-2 homologs. Autophagy 4, 268–272 (2008).

17. Liang, Q. et al. Kaposi’s sarcoma-associated herpesvirus K7 modulates Rubicon-mediated inhibition of autophagosome maturation. J. Virol. 87, 12499–12503 (2013).

18. Orvedahl, A. et al. HSV-1 ICP34.5 confers neurovirulence by targeting the Beclin 1 autophagy protein. Cell Host Microbe 1, 23–35 (2007).

19. Klionsky, D. J. et al. Guidelines for the use and interpretation of assays for monitoring autophagy (3rd edition). Autophagy 12, 1–222 (2016).

20. Gassen, N. C. et al. Analysis of SARS-CoV-2-controlled autophagy reveals spermidine, MK-2206, and niclosamide as putative antiviral therapeutics. bioRxiv 2020.04.15.997254 (2020) doi:10.1101/2020.04.15.997254.

21. Kimura, S., Noda, T. & Yoshimori, T. Dissection of the autophagosome maturation process by a novel reporter protein, tandem fluorescent-tagged LC3. Autophagy 3, 452–460 (2007).

22. Li, J. et al. Virus-Host Interactome and Proteomic Survey Reveal Potential Virulence Factors Influencing SARS-CoV-2 Pathogenesis. Med N. Y. N (2020) doi:10.1016/j.medj.2020.07.002.

23. Gordon, D. E. et al. Comparative host-coronavirus protein interaction networks reveal pan-viral disease mechanisms. Science (2020) doi:10.1126/science.abe9403.

24. Gordon, D. E. et al. A SARS-CoV-2 protein interaction map reveals targets for drug repurposing. Nature 583, 459–468 (2020).

25. Liang, C. et al. Autophagic and tumour suppressor activity of a novel Beclin1-binding protein UVRAG. Nat. Cell Biol. 8, 688–698 (2006).

26. Liang, C. et al. Beclin1-binding UVRAG targets the class C Vps complex to coordinate autophagosome maturation and endocytic trafficking. Nat. Cell Biol. 10, 776–787 (2008).

27. Matsunaga, K. et al. Two Beclin 1-binding proteins, Atg14L and Rubicon, reciprocally regulate autophagy at different stages. Nat. Cell Biol. 11, 385–396 (2009).

28. Zhong, Y. et al. Distinct regulation of autophagic activity by Atg14L and Rubicon associated with Beclin 1- phosphatidylinositol-3-kinase complex. Nat. Cell Biol. 11, 468–476 (2009).

29. Zhong, N. S. et al. Epidemiology and cause of severe acute respiratory syndrome (SARS) in Guangdong, People’s Republic of China, in February, 2003. The Lancet 362, 1353–1358 (2003).

30. Brest, P., Benzaquen, J., Klionsky, D. J., Hofman, P. & Mograbi, B. Open questions for harnessing autophagy-modulating drugs in the SARS-CoV-2 war: hope or hype? Autophagy 0, 1–4 (2020).

31. Mijaljica, D. & Klionsky, D. J. Autophagy/virophagy: a “disposal strategy” to combat COVID-19. Autophagy 0, 1–2 (2020).

32. Wolff, G. et al. A molecular pore spans the double membrane of the coronavirus replication organelle. Science (2020) doi:10.1126/science.abd3629.

33. Maisonnasse, P. et al. Hydroxychloroquine use against SARS-CoV-2 infection in non-human primates. Nature 1–8 (2020) doi:10.1038/s41586-020-2558-4.

34. Gao, J., Tian, Z. & Yang, X. Breakthrough: Chloroquine phosphate has shown apparent efficacy in treatment of COVID-19 associated pneumonia in clinical studies. Biosci. Trends 14, 72–73 (2020).

35. Colson, P., Rolain, J.-M., Lagier, J.-C., Brouqui, P. & Raoult, D. Chloroquine and hydroxychloroquine as available weapons to fight COVID-19. Int. J. Antimicrob. Agents 55, 105932 (2020).

36. Savarino, A., Boelaert, J. R., Cassone, A., Majori, G. & Cauda, R. Effects of chloroquine on viral infections: an old drug against today’s diseases. Lancet Infect. Dis. 3, 722–727 (2003).

37. Devaux, C. A., Rolain, J.-M., Colson, P. & Raoult, D. New insights on the antiviral effects of chloroquine against coronavirus: what to expect for COVID-19? Int. J. Antimicrob. Agents 55, 105938 (2020).

38. Hoffmann, M. et al. Chloroquine does not inhibit infection of human lung cells with SARS-CoV-2. Nature 1–5 (2020) doi:10.1038/s41586-020-2575-3.

39. Liang, Q. et al. Crosstalk between the cGAS DNA sensor and Beclin-1 autophagy protein shapes innate antimicrobial immune responses. Cell Host Microbe 15, 228–238 (2014).

